# Measuring the temporal quality of a biodiversity database

**DOI:** 10.1101/2021.07.27.453997

**Authors:** Les G Underhill

**Affiliations:** Department of Biological Sciences, University of Cape Town, Rondebosch, South Africa

## Abstract

In the context of climate change it is important to keep biodiversity databases up-to-date. This priority generates the need for a metric to assess the concept of up-to-dateness. The objective of this paper is to devise a measure of up-to-dateness for atlas-type biodiversity data. The data input into the algorithm consists of the species, date and grid cell allocation of all available records for a taxon in a region. First, for each grid cell in a region, the median of the date of the most recent record of each species is calculated. Secondly, the median of the median dates for each grid cell yields an overall measure of up-to-dateness. The performance of this algorithm is investigated in relation to databases for six taxa in southern Africa. In June 2021, the up-to-dateness of the databases varied from 41 years for the reptile database to two years for the bird database. The quality of a biodiversity database is a multidimensional concept; up-to-dateness is only one of several dimensions. The paper identifies a need to quantify the rate at which the “value” of a record decays as evidence that a species still occurs at a locality, and suggests an experimental process for doing this. The use of the up-to-dateness index to motivate citizen scientists is discussed.

## Introduction

Sutherland et al. (2015) developed a set of 10 priorities for biological recording, with a focus on monitoring biodiversity through citizen science. This paper develops an 11th priority which, though not made explicit by Sutherland et al. (2015), was probably in mind. In the context of climate change, it is necessary to articulate the conscious and continuous need to keep biodiversity databases up-to-date; a corollary to this priority is the need for a metric to assess the concept of up-to-dateness. The objective of this paper is to meet this need to quantify up-to-dateness, and thus provide one measure of the temporal quality of a biological database.

The context of this paper is the ongoing collection of “atlas”-type data for a taxon, used to generate on-demand, up-to-date distribution map for any species in that taxon. Such maps are of fundamental value in establishing conservation priorities (Underhill & Gibbons 2002, Harrison et al. 2008). For example, the Second Southern African Bird Atlas Project began in 2007 to serve as a five-year “snapshot” of bird distributions, as had been the objective of the initial bird atlas. Instead, the project remains ongoing in 2021, having morphed from the aim of capturing a “snapshot” to recording a “video” of changing distributions (Harrison et al. 2008, Underhill 2016, Underhill et al. 2017). The key idea is that up-to-date distribution maps should be built on recent data. Representing the occurrence of a species at a location with an “old” record is unsatisfactory, and even misleading.

Therefore, the focus of this paper is to devise an algorithm which, in some way, measures the decay in temporal quality of biodiversity data. The algorithm attempts to answer the question: “how up-to-date is this collection of biodiversity records?”. The algorithm is illustrated using the South African component of the Biodiversity and Development Institute’s Virtual Museum database, which includes South Africa, Lesotho and eSwatini; however, the method is readily adapted to other contexts.

## Methods

### Algorithm

A biodiversity record is comprised of three components: species, place and date. The record provides evidence that a particular species was recorded at the place on the date. The algorithm proposed here requires each data point to be allocated to a “grid cell”. For the Virtual Museum, a 15-minute geographical grid is used, generating what are popularly known as quarter-degree grid cells (even though there are 16 of them in a one-degree grid cell). For the African Bird Atlas Project, a five-minute grid is in use, generating grid cells known as pentads (Underhill 2016). North of c. 40°N, geographical grid cells are no longer viable, because the east-west distance is far smaller than the north-south distance. The convention then is to use grid cells measured in kilometres, as is done in the bird atlas projects of Europe (e.g. Hagemeijer & Blair 1997). In Britain, early projects made use of “counties” and “vice-counties”; the proposed method works with any spatial units, provided they are standardized through time.

For each record, the algorithm needs the three components: the date, the species, and the grid cell to which it belongs. The algorithm itself has four steps:

1. For each grid cell, find the most recent date for each species;
2. Find the median of these dates, one per species (this date provides a measure of the temporal quality of the data for the grid cell);
3. Find the median dates for all grid cells in the region under consideration;
4. Calculate the median of the grid cell medians (this date, the median of the medians, is defined to be the temporal quality of the data in the region, i.e. up-to-dateness).

The algorithm is simple and transparent. At Step 2, the median date calculated for a grid cell is the halfway point for the most recent dates for the species recorded in that grid cell. At Step 4, the halfway point for all the grid cells in the region is calculated. The median dates of half of the grid cells fall before this date, and half fall later.

The algorithm can be applied not only to the current database, but also to historic databases created by only considering records submitted prior to a chosen date. The crucial statistic then becomes the time interval, expressed in months or years, between the median of the medians and the chosen date. If this interval is shortening through time, then the temporal quality of the database is improving, and *vice versa*. The trend in this statistic through time is a powerful communication tool.

### Application

The algorithm was applied to the databases of six of the projects within the Virtual Museum. The Virtual Museum is a citizen science initiative which assembles photographic records for a series of taxa. It was originally developed for citizen scientists to contribute to reptile and butterfly atlases in South Africa (Bates et al. 2014, Mecenero et al. 2013), and gradually expanded to cover a selection of other taxa. The databases for reptiles, butterflies, dragonflies and damselflies (Underhill et al. 2016), lacewings (Underhill et al. 2019), birds and frogs were evaluated. The algorithm used all records up to 22 June 2021. The “popular” names for the six databases are included in Table 1.

**Table 1.**
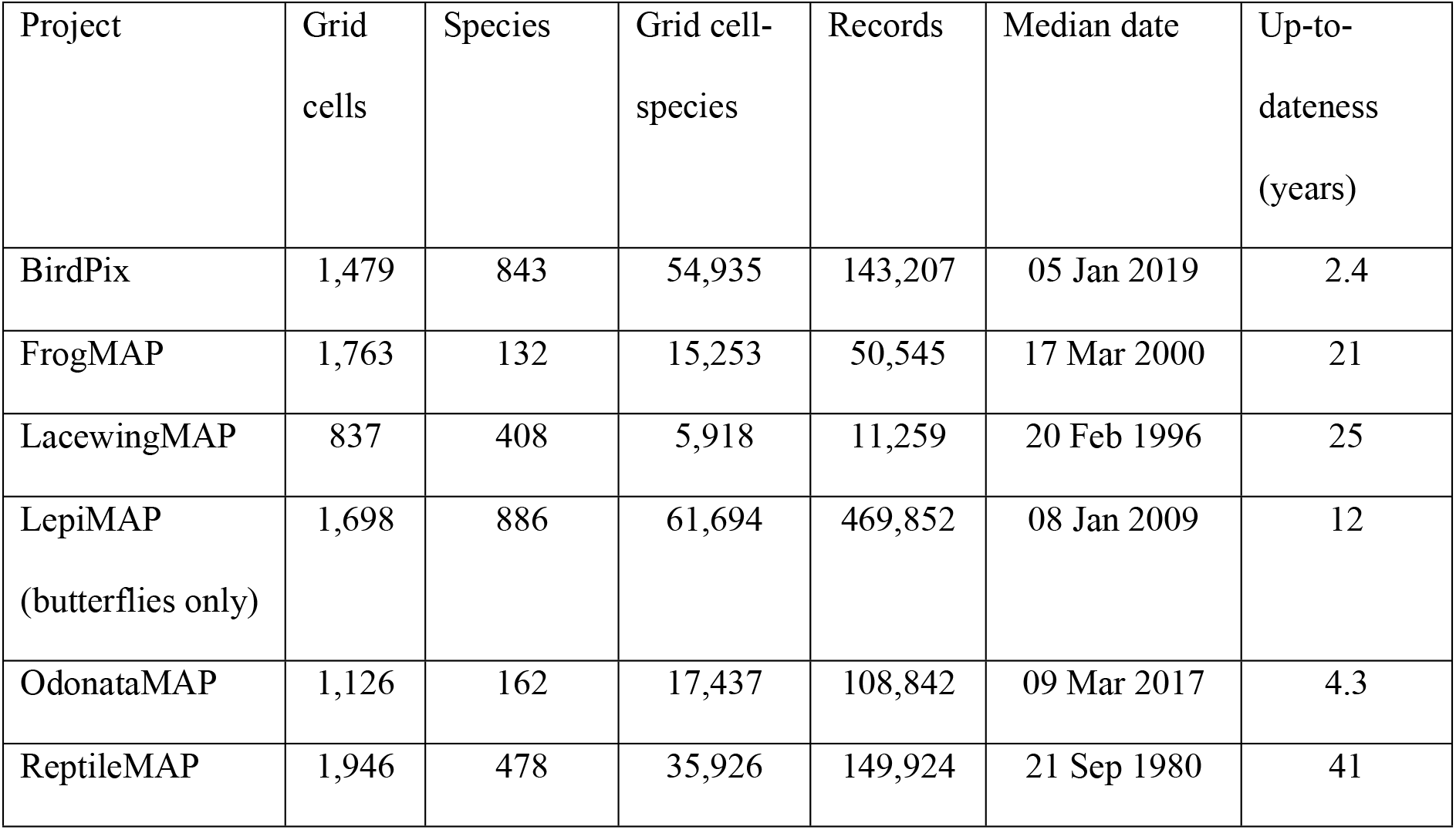
Summary statistics, including up-to-dateness, for six projects within the Virtual Museum, in the 2,027 quarter degree grid cells in South Africa, Lesotho and eSwatini. Calculations were performed on 25 June 2021, and the up-to-dateness is calculated as the difference between the median date and 25 June 2021.

| Project | Grid cells | Species | Grid cell-species | Records | Median date | Up-to-dateness (years) |
| --- | --- | --- | --- | --- | --- | --- |
| BirdPix | 1,479 | 843 | 54,935 | 143,207 | 05 Jan 2019 | 2.4 |
| FrogMAP | 1,763 | 132 | 15,253 | 50,545 | 17 Mar 2000 | 21 |
| LacewingMAP | 837 | 408 | 5,918 | 11,259 | 20 Feb 1996 | 25 |
| LepiMAP<br>(butterflies only) | 1,698 | 886 | 61,694 | 469,852 | 08 Jan 2009 | 12 |
| OdonataMAP | 1,126 | 162 | 17,437 | 108,842 | 09 Mar 2017 | 4.3 |
| ReptileMAP | 1,946 | 478 | 35,926 | 149,924 | 21 Sep 1980 | 41 |

To illustrate the interpretation of the pattern in trends through time, the algorithm was also applied to dragonflies and damselflies records (OdonataMAP) on an annual basis over the decade since the project commenced in September (Underhill et al. 2016). To assess whether the rate of renewal of old records resulted in distribution maps becoming more or less up-to-date through time, the algorithm was applied to observations uploaded by the ends of the six calendar years from 2015 to 2020.

## Results

Of the six Virtual Museum projects for which up-to-dateness was evaluated in June 2021, it is clear that the distribution maps produced using the ReptileMAP data cannot be considered “up-to-date”. For half of the grid cells, the median date of the most recent record was 41 years prior to the date of the analysis. Similarly, the LacewingMAP database was 25 years out of date, the FrogMAP database 21 years out of date, the LepiMAP database 12 years out of date, and the OdonataMAP database 4.3 years out of date. The BirdPix database was classified as 2.4 years, out of date (Table 1).

For OdonataMAP, the pattern of up-to-dateness reflects the typical behaviour anticipated as a project evolves (Table 2). Over the first years to the end of 2013, coverage expanded rapidly, and up-to-dateness remained constant. Thereafter, without an incentive to “refresh” records of a species in a grid cell the up-to-dateness steadily moves backwards. Since 2016, a large number of historical museum specimen records were included in the analysis, causing a 24-month retrogression in up-to-dateness. In the subsequent four years from the end of 2016 to the end of 2020, the up-to-dateness of the OdonataMAP database had slipped by four months, from 46 months at the end of 2016 to 50 months at the end of 2020 (Table 2). At the end of 2020, for half of the 1098 grid cells visited, the median date of the most recent observation of a species was after 10 November 2016, and before this date for the other half.

**Table 2.** Summary statistics, including up-to-dateness, for the OdonataMAP project of the Virtual Museum in the 2,027 quarter degree grid cells of South Africa, Lesotho and eSwatini. These statistics are calculated from the database using the records that had been uploaded by the end of each of the six calendar years from 2011 to 2020.

| Period | Grid cells | Species | Grid cell-species | Cumulative records | Median date | Up-to-dateness (months) |
| --- | --- | --- | --- | --- | --- | --- |
| Up to 2011 | 99 | 81 | 466 | 534 | 04 Dec 2010 | 13 |
| Up to 2012 | 242 | 118 | 1,344 | 2,332 | 12 Dec 2011 | 13 |
| Up to 2013 | 395 | 133 | 2,863 | 6,001 | 06 Jan 2013 | 12 |
| Up to 2014 | 529 | 145 | 4,438 | 9,591 | 05 May 2013 | 20 |
| Up to 2015 | 679 | 147 | 5,954 | 14,994 | 22 Feb 2014 | 22 |
| Up to 2016 | 852 | 159 | 11,620 | 36,900 | 17 Mar 2013 | 46 |
| Up to 2017 | 925 | 160 | 13,227 | 48,441 | 09 Feb 2014 | 47 |
| Up to 2018 | 1,008 | 160 | 14,656 | 65,759 | 06 Feb 2015 | 47 |
| Up to 2019 | 1,062 | 162 | 15,739 | 82,497 | 14 Dec 2015 | 49 |
| Up to 2020 | 1,098 | 162 | 16,556 | 98,505 | 10 Nov 2016 | 50 |

## Discussion

### Multiple measures of quality

When applied to a biodiversity database, the term “data quality” is frequently used qualitatively rather than quantitatively. For example, Wetzel et al. (2018) repeatedly used the word “quality” in describing the needs of biodiversity data in Europe, but never clearly defined the term. There is a need to quantify the concept of “quality.” In this context, “quality” is a multi-dimensional concept; there is no single measure of quality.

Up-to-dateness, as presented here, is only one such measure of quality. It must be supported by supplementary information which captures other dimensions of quality, i.e. the total number of records within the region, the number of species represented, the number of grid cells with records, and the number of grid cell-species records (the sum of the number of species recorded in each grid cell). A further family of data quality refer to taxonomic issues: the up-to-dateness of the taxonomy used for the database, and the percentage of records not correctly identified to species level.

There are several qualitative dimensions implicit in the concept of biodiversity data quality. The most frequently employed relates to gaps in coverage, i.e. spatial quality. Data are of poor quality if they display gaps. These are either geographical gaps, areas for which little or no biodiversity data exists for an entire taxon; or gaps in the range map for an individual species, i.e. places where the species probably occurs, but has not yet been recorded (false negatives). A second dimension is yet another measure of temporal quality, which is generally interpreted to mean that there are data for an extended time period, usually measured in years or decades. In this context, biodiversity data for a region are said to have poor temporal quality if they lack historical records, making it difficult to examine trends through time.

For example, with data from 1,946 of 2,027 grid cells, ReptileMAP has the best spatial coverage (96%) of the six projects (Table 1). More nuanced measures of spatial quality would need to 1) account of whether gaps in coverage tend to be in adjacent or scattered grid cells, and 2) estimate the percentage of false negatives in the database. For 2), determining the number which goes into the numerator when estimating this percentage is straightforward; it is the sum of the numbers of species in each grid cell, 35,926 in the case of ReptileMAP (Table 1). The denominator for the percentage requires an estimate of the number of grid cells in the range of each species, a quantity which the atlas project aims to determine. The number 35,926 is the total of the total number of grid cells in which the 478 reptile species have been recorded, and hence shaded in the distribution maps (Table 1).

Table 2 details improvements of other measures of quality in OdonataMAP data through time. A set of distribution maps for 147 species made at the end of 2015 would have been based on 14,994 records from 679 grid cells. In these maps, the total number of grid cells which would have shown a species as present would have been 5,954 (Table 2). One year later, after the inclusion of the historical data, maps for 159 species would have been based on 36,900 records from 852 grid cells, and the total number of shaded grid cells would have almost doubled to 11,620 (Table 2). The improved spatial coverage was obtained by sacrificing up-to-dateness (Table 2). By June 2021, the set of 162 maps including the additional records collected from 2017 onwards would have had a total of 17,437 grid cells showing a species as present (Table 1). This is an increase of 50% since the end of 2016, providing another useful measure of the improvement of the spatial quality of the OdonataMAP database in South Africa, Lesotho and eSwatini in 4.5 years.

Thus quality, in this context, is a multidimensional concept. Temporal quality, as developed here, is only one dimension in describing the value of the database.

### An alternative approach to measure the temporal decay in quality

A biodiversity record provides evidence that a particular species was recorded at the place on the date. However, biodiversity literature seems to pay little or no attention to the reality that the value of a record steadily decreases as its date of occurrence recedes into the past. Aging records slowly but steadily become less and less valuable as evidence that a species still occurs at a given site.

In this paper, the concept of temporal quality of a biodiversity database is introduced, rooted in the notion that data quality decays over time. However, the shape of the function which describes the decay in value through time remains unknown, and needs to be quantified.

Here is an experimental approach to achieve this. Consider a substantially-sized sample (say *n* = 1000) of correctly georeferenced biodiversity records with accurate dates. The sample needs to be stratified geographically and by date, possibly from one year ago to 100 years ago. Revisit the sites at which the records were made on the calendar date of the original record, and search for the species. Three outcomes are possible: (1) the species was recorded, and still occurs at the site, (2) there is still suitable habitat for the species, but it was not found, and (3) the site has been transformed to such an extent that the species almost certainly no longer occurs. The data analysis should aim to develop a function which describes the average rate of decay in value of biodiversity records, i.e. to estimate the “half-life” of a record. It is likely that these decay functions vary between species. One objective, probably unattainable, would be to establish a gold standard definition for the age at which a record no longer provides suitable evidence that a species still occurs at a locality. Distribution maps for a species could then exclude records older than this date, or plot them differently to indicate regions in which the species has either actually gone locally extinct, or where further search effort is needed to refresh evidence of its presence. Alternatively, distribution maps could weight records by their age, so that older records have smaller weights than newer ones.

The decay function also opens up new possibilities of developing a more nuanced measure of up-to-dateness than the one developed here; for instance, using the last recorded date for each species in a grid cell, one could apply the decay function to estimate the remaining evidential value of the record, and calculate appropriate summary statistics.

### Use of up-to-dateness to motivate citizen scientists

As demonstrated above for OdonataMAP (Table 2), the algorithm is not only applicable to the current database. It can also be applied to generate trends from historical databases, created by only considering records submitted up to a certain point in time. When applied in this way, these trends may serve as motivational guidance for project leaders and citizen scientists.

For instance: “In this era of rapid development and climate change, any record of a species at a locality which is more than two years old needs to be refreshed with a new one.” The awareness of last recorded dates for the species in a grid cell is in itself a powerful motivational force for citizen scientists. There is a real challenge in seeking to detect the species that have the most-out-of-date last recorded dates in a grid cell.

The background to the six projects of Table 1 provides insights into the extent of their out-of-dateness. This information must be considered in communications with the citizen scientists participating in each project. Three of these projects (FrogMAP, LepiMAP and ReptileMAP) are continuations of citizen science projects which produced published atlases for frogs (Minter et al. 2004), butterflies (Mecenero et al. 2013) and reptiles (Bates et al. 2014) respectively. In each case, the foundational data for the project were drawn from historical databases in museums, other non-museum specimen collections, and literature. The explicit objective of the citizen science fieldwork, which took place over five-year periods, was to fill in distribution gaps with a goal of targeting false negatives, grid cells where the species likely occurred but had not yet been recorded. This mentality of “filling in the gaps,” has prevailed in the continuation of these projects within the Virtual Museum. Thus, many observers do not upload records to the Virtual Museum if the species has already been recorded in the grid cell. The consequence is that the up-to-dateness of the databases slowly deteriorates. Changing this outlook has proved challenging.

OdonataMAP is a relatively new project (Underhill et al. 2016), and the cohort of citizen scientists involved are relatively strongly attuned to the importance of repeated submission of the same species from the same locality. To a large measure, this is because the project includes a focus on generating data from which the phenology of adult dragonfly and damselfly occurrence may be estimated. A by-product of this approach is that overall up-to-dateness of the database deteriorated by only four months over four years (Table 2).

The foundational data for LacewingMAP were developed by Mervyn Mansell over a career; they contain the global specimen database for the Neuroptera and Megaloptera, with records dating back to the nineteenth century (Underhill et al. 2019). The citizen science database is built on this platform. Contributions are opportunistic; no citizen scientist has a primary interest in this taxon (Underhill et al. 2019). The rate of record submission is thus relatively slow (but far faster than the rate of specimen acquisition in the decades when museum collections were growing fastest (Underhill et al. 2019)). Thus, the fact that the database is 25 years out of date is not surprising (Table 2).

In contrast, although the BirdPix project of the Virtual Museum was started in 2012, active promotion of the project commenced in 2017. By then a large number of grid cells had received a small number of records each. Many citizen scientists have uploaded series of historical photos taken in the early days of digital photography, or have even scanned and submitted slides; this has effectively pushed the median of medians further into the past. For the other projects within the Virtual Museum discussed here, most historical image collections (for example of reptiles and butterflies) were uploaded during the formal atlas periods for each taxon. However, in spite of these challenges, the BirdPix database was 2.4 years out of date in June 2021.

The continuous presentation of up-to-dateness, preferably in graphical form, provides an incentive to citizen scientists to keep the database up-to-date. This is an example of the application of gamification to motivate data collection by citizen scientists (Ainsley & Underhill 2017). Gamification is the development of built-in strategies to encourage project participation (in particular, it does not mean turning data-collection into a “game”) (Ainsley & Underhill 2017).

For a citizen scientist to contribute towards bringing the median date for a grid cell closer to the present, a straightforward strategy is to increase the number of species for that grid cell, especially if it is still relatively easy to add species; each added species then has a current date and moves the median for the grid cell forward. Similarly, to bring the median of medians closer to the present would require undertaking fieldwork in grid cells which have no data. This increases the number of grid cells with median dates; the new median dates are current, and move the median of medians forward. Both strategies not only improve up-to-dateness, they also positively impact other measures of completeness for the database (i.e. spatial coverage).

### Local extinction

Clearly, if a species goes extinct in a grid cell, its last recorded date can no longer be updated. At some point, decisions about local extinction need to be made, so that a species can be removed from the list over which the median for the grid cell is calculated. Mechanisms to do this need to be devised, but will almost certainly be a combination of quantitative and qualitative arguments. Species declared locally extinct need to remain on the species list, and flagged appropriately. The awareness among citizen scientists of local extinctions has the potential to lead to civic awareness so that the impact of the project transcends science into grassroots action and policy making (Loos et al. 2015).

### Caveats to the measure of up-to-dateness

The measure of up-to-dateness described here is conditional on two factors: first, it only takes account of grid cells for which data are available; and second, it only takes account of species which have already been recorded in the grid cell. An absolute measure of up-to-dateness would also account for the grid cells lacking data, and include an estimate of the number of species in each grid cell.

The choice of the conditional measure was deliberate. Primarily, it is designed to measure the up-to-dateness of the data already collected; other aspects of database quality can be evaluated using other statistics, such as spatial coverage measures. Secondly, the choice of conditional measure facilitates the use of gamification, as described above.

## Acknowledgements

Itxaso Quintana, Magda Remisiewicz, Karis Daniel, Johan van Rooyen, Greg Distiller and David Thomson made important comments on earlier drafts. The paper owes its existence to the thousands of citizen scientists who have participated in the Virtual Museum.

## Notes

### Competing Interest Statement

The authors have declared no competing interest.

